# Prospective Motion Correction and Automatic Segmentation of Penetrating Arteries in Phase Contrast MRI at 7 T

**DOI:** 10.1101/2022.01.20.477093

**Authors:** Julia Moore, Jordan Jimenez, Weili Lin, William Powers, Xiaopeng Zong

**Author notes:** Corresponding Author: Xiaopeng Zong, Department of Radiology and Biomedical Research Imaging Center, University of North Carolina at Chapel Hill, CB#7515 Chapel Hill, NC, 27599.

## Abstract

**Purpose:** To develop a prospective motion correction (MC) method for phase contrast (PC) MRI of penetrating arteries (PA) in centrum semiovale at 7 T and evaluate its performance using automatic PA segmentation.

**Methods:** Head motion was monitored and corrected during the scan based on fat navigator images. Two convolutional neural networks (CNN) were developed to automatically segment PAs and exclude surface vessels. Real-life scans with MC and without MC (NoMC) were performed to evaluate the MC performance. Motion score was calculated from the range of translational and rotational motion parameters. MC vs NoMC pairs were divided according to their score differences into groups with similar, less, or more motions during MC. Data reacquisition was also performed to evaluate whether it can further improve PA visualization.

**Results:** In the group with similar motion, more PA counts (N_PA_) were obtained with MC in 9 (60%) cases, significantly more than the number of cases (1) with less PAs (p = 0.011; binomial test). In the group with less motion during MC, MC images had more or similar NPA in all cases, while in the group with more motion during MC, the numbers of cases with less and more NPA during MC were not significantly different (3 vs 0). Data reacquisition did not further increase N_PA_. CNNs had higher sensitivity (0.85) and accuracy (Dice coefficient 0.85) of detecting PAs than a threshold based method.

**Conclusions:** Prospective MC and CNN based segmentation improved the visualization and delineation of PAs in PC MRI at 7 T.

## 1. INTRODUCTION

Cerebral small vessel disease (SVD) is a condition postulated to have multiple different etiologies. It may result from a cascade of events, starting with endothelial dysfunction in the penetrating arteries (PAs) [1]. Other causes include the thickening of the arterial media due to lipohyalinosis or obstruction of the origins of PAs by parent artery intimal plaques, which lead to brain ischemia resulting in deep small infarcts and leakage of fluid causing edema and later gliosis in white matter tracts [2]. Additionally, inflammation might also be involved in the development of PA leakage and endothelial dysfunction, although the causal relationships are yet to be established [3]. Cumulatively, SVD might lead to substantial cognitive [4–6], psychiatric[7], and physical disabilities [8, 9].

Phase contrast (PC) MRI is a common method for imaging arteries and can visualize a large number of PAs (27 – 91) in centrum semiovale at 7 T [10, 11]. One major concern that arises during imaging is the susceptibility to motion which reduces PA visibility and might introduce artificial group differences due to different motion tendencies between the participants. As the PC MRI only covers a single slice, prospective motion correction (MC) is needed to ensure consistent imaging location in the presence of head motion. In this study, we will evaluate the improvements of PA visualization by prospective MC based on motion parameters derived from whole-brain fat navigator (FatNav) images [12].

Accurate segmentation of penetrating arteries is a key step in objective evaluation of improved PA visualization by MC and in future studies delineating the roles of PAs in small vessel disease. The existing segmentation method relied on thresholding of the phase and magnitude image intensities which only controls for false positives rate, but might suffer from a high false negative rate due to the low contrast to noise ratio of some PAs in the images [10, 11]. Here, we evaluate the performance of a two-dimensional multi-channel multi-scale encoder decoder network (M2EDN) for segmenting the PAs, which was adopted from a three-dimensional M2EDN originally developed for perivascular spaces segmentation [13]. Furthermore, since some arteries on cortical surface mimic penetrating arteries, a white matter (WM) mask can be applied to remove such false PAs and further improve segmentation accuracy. To this end, we also developed a 3D U-NET based method to obtain the WM mask, utilizing 3D T2-weighted (T2w) images that were acquired during the same imaging session [14].

## 2. METHODS

### 2.1 Subjects

This study was approved by the Institutional Review Board of the University of North Carolina at Chapel Hill. Written informed consents were obtained from all subjects before the scan. Two separate experiments were performed: Experiment 1 (Exp 1) acquired images used for training the convolutional neural networks (CNN) parameters and Experiment 2 (Exp 2) were carried out to evaluate the performance of MC. 39 healthy volunteers (aged 21 – 55 years, 28 females) were included in Exp 1 and 21 patients with diabetes and 13 healthy controls (aged 37 – 70 years, 19 females) were included in Exp 2.

### 2.2 Data acquisition

All images were acquired on a 7T MRI scanner (Siemens Healthineer, Erlangen, Germany). In Exp 1, a Nova 32-channel receiver and 8-channel transmitter head coil (Nova Medical, Wilmington, MA, USA) was used. No radio frequency magnetic field (B_1_) shimming was performed. In Exp 2, the images were acquired using a Nova 32-channel receiver and single-channel volume transmitter coil. The subjects were asked to keep their heads still during all scans.

A 3D variable flip angle turbo spin echo (TSE) sequence was used to acquire T2w images for WM segmentation [15, 16]. A FatNav image [12] was acquired within each repetition time (TR) to monitor head motion. In Exp 1, the FatNav images were not utilized to correct motion in the present study, although they had been utilized to perform retrospective motion correction in a separate study [17]. In Exp 2, the FatNav images were used to estimate motion parameters relative to the first FatNav image which were then used to adjust the imaging FOV for the next TR.

Then, a single slice PC MRI was performed to image PAs in centrum semiovale. The slice was positioned 15 mm above the corpus callosum and parallel to the anterior commissure – posterior commissure line. No FatNav was acquired for the PC MRI scans in Exp 1. In Exp 2, the FatNav and PC MRI data were acquired alternatively as shown in the sequence diagram in Fig. 1. After acquiring and reconstructing each FatNav, motion parameters were estimated using vendor provided module (MOCO functor) and the PC MRI slice position and readout, phase encoding, and slice normal directions were adjusted accordingly. As the slice was prescribed based on the TSE images, to correct for potential motion between the TSE and PC MRI scans, the FatNavs in PC MRI were registered to the first FatNav from the TSE-MRI scan.

**Figure 1:**
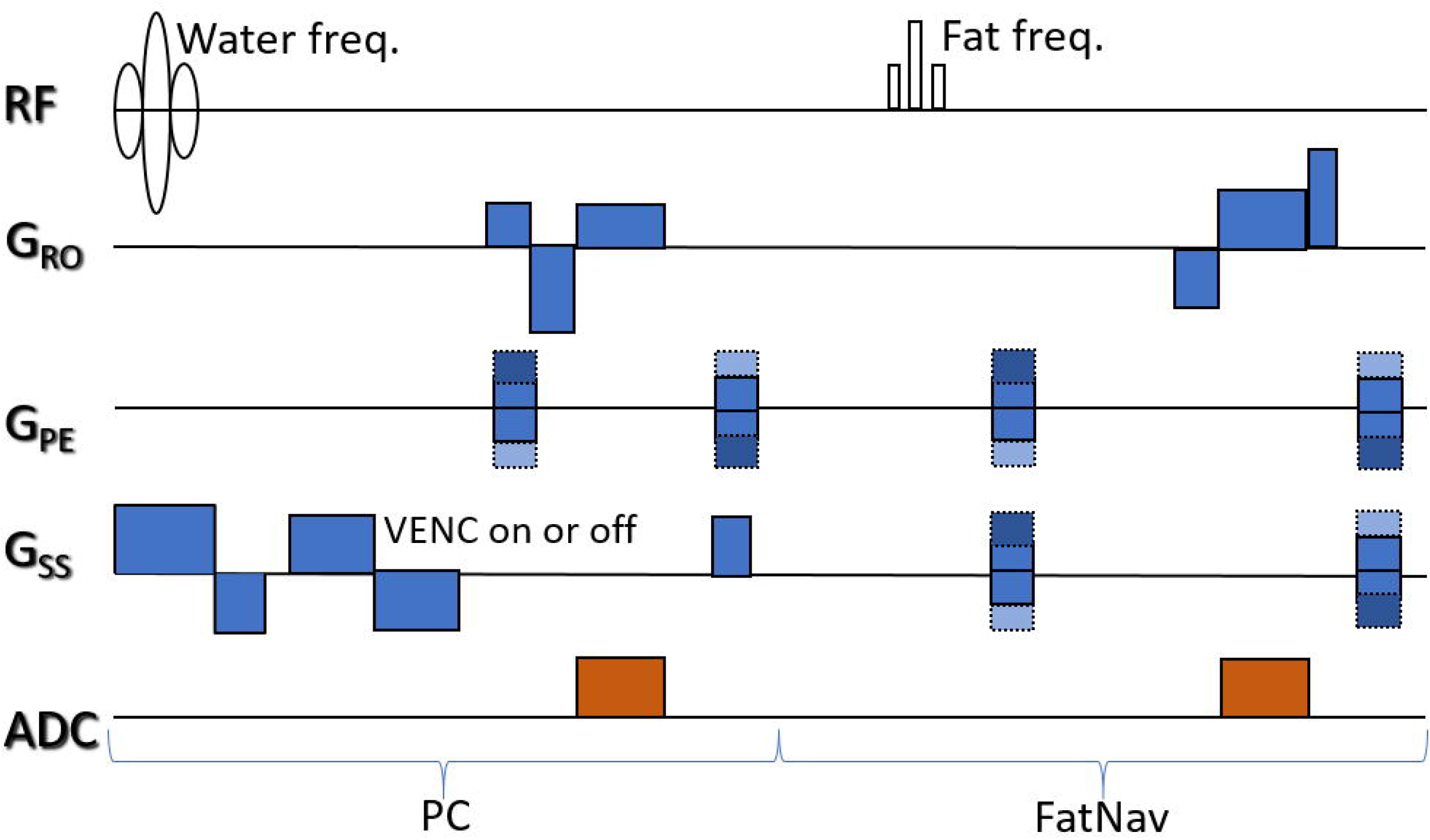
Pulse sequence of the phase contrast-MRI sequence with FatNav-based motion correction.

Because it took 4.68 s to acquire a complete FatNav in PC MRI and an additional 2.1 s to reconstruct the image, motion occurred due this period cannot be corrected. Therefore, reacquiring k-space data during periods with large motion may further improve image quality. For this purpose, k space data were divided by the times when the motion parameter feedbacks were received into blocks. Denote the motion parameters received at the beginning and end of data block *i* (*D_i_*) as *M*_i-1_ and *M_i_*, corresponding FatNav images as *F*_i-1_ and *F*_i_, and the relative translation and rotation between *F*_i-1_ and *F*_i_ as *T*_i_ and *R*_i_ which can be calculated from *M*_i-1_ and *M*_i_. *D*_i_ was reacquired if *T*_i-1_, *T*_i_, or *T*_i+1_ was greater than 0.1 mm or *R*_i-1_, *R*_i_, or *R*_i+1_ was greater than 0.1°. The requirements ensured that the motion was below the specified thresholds during the *F*_i-1_, *F*_i_, and *D*_i_ acquisition periods. The reacquired data was in the same temporal order as the original data. After the completion of data reacquisition, the same criterion was used to determine whether the required data needed to be reacquired one more time. The scan stopped until no further reacquisition was needed or the total reacquisition time exceeded the original scan time.

Three (n = 32) or two (n = 1) PC MRI scans with MC were performed in each subject. Data reacquisition was carried out in all scans in 18 subjects but only in the last scan in 15 subjects. For comparison, a fourth scan without MC (NoMC) was performed in 14 subjects where the slice position was not updated while the FatNavs were still acquired for monitoring motion. Furthermore, to study whether the addition of the FatNav images would impact the quality of PC MRI images, the PC MRI with MC and the conventional PC MRI scan without the FatNav module were acquired in an additional subject, while the subject kept the head still.

At the beginning of each sequence that contained FatNav, a navigator image that included fully sampled GRAPPA calibration data at the k-space center was acquired. The parameters for all the sequences are provided in Supporting Information Table S1.

### 2.3 Segmentation

#### M2EDN for PA segmentation

The network architecture for PA segmentation is the same as that proposed in [13] except for two modifications. First, the phase and magnitude images were used as network input channels instead of the original and preprocessed images. Second, the convolution and pooling layers were changed from 3-dimensional to 2-dimensional since the input images were twodimensional. In addition, due to the smaller memory size required for 2D images, the entire images were fed into the network without separating them into patches.

#### U-NET for WM segmentation

The network consists of 4 steps in the contracting path and another 4 steps in the expansive path. The contracting path applies two 3×3×3 convolutional layers in each step. In between these convolutional layers, we incorporated a dropout layer of 0.1, 0.2, or 0.3. And specific to the contracting path, different spatial scales were connected by 2×2×2 max-pooling layers with a stride of 2 [18]. We started with an initial patch size of 128×128×128 and 16 channels. Channel number doubled with each step in the contracting path (32, 64, 128, 256) but patch size was halved with each step (64×64×64, 32×32×32, 16×16×16, 8×8×8...). The same pattern exists in the expansive path back to 16 channels and 128×128×128 patch size. The expansive path upsamples the image back to its original size using transposed convolutions. Once the image is upsampled, it is combined with the image at the same spatial scale from the contracting path to get a more accurate result. Lastly, a 1×1 convolutional layer is applied to produce the output with a 5 channels, which is followed by a softmax layer to produce the probabilities of the background plus four tissue types, including thalamus, basal ganglia, WM, and midbrain, the same ROIs as in our previous study [19, 20]. The architecture of the U-NET can be found in Supporting Information Fig. S1.

#### Ground truth

Ground truth (GT) masks for training the networks were manually drawn in ITK-Snap and custom made Matlab (MathWorks, Natick, MA, USA) programs. The four tissue ROIs were defined in 10 TSE images from Exp 1. PAs masks were defined as hyperintensive voxels in the phase image inside WM on 61 PC MRI images from Exp 1. To control for false positive voxels, each hyperintensive cluster had to overlap with a hyperintensive cluster on the corresponding magnitude image.

#### Training and testing

Both networks were implemented using Keras and trained using the ground truth masks defined above. The image – GT mask pairs were divided into 10 groups. Nine groups were selected for training and the remaining one for testing the model performance. To enhance the training dataset, new training data were generated by the following procedures: 1) flip the T2w image – GT mask pairs in the L-R direction, 2) rotate the pairs by 10 and −10 degrees around the I-S and L-R axes. 3) flip the PC MRI image – GT mask pairs in the L-R direction. The training was performed for 40 Epochs with batch sizes of 4 and 2 for U-NET and M2EDN. Convergence of the training was confirmed by plotting the sensitivity (SEN), positive predictive value (PPV), and dice similarity coefficient (DSC) versus the number of epochs. For U-NET, the Adam optimizer was employed with a learning rate of 0.0001 and categorical cross entropy loss function For M2EDN, the same optimizer was employed but with a learning rate of 0.001 and the loss function was the same as defined in [13].

Using the group that were left out for testing, we numerically analyzed segmentation performances of our networks using SEN, PPV, and DSC as defined below:

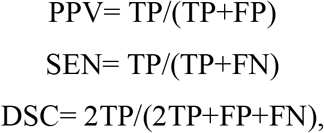

where TP, FP, and FN stand for the number of true positive, false positive, and false negative voxels, respectively, using the manually drawn masks as ground truth. Additionally, since the true size of PAs are less than 1 voxel in the PC MRI images and the main goal of the segmentation algorithm is to simply detect visible PAs, but not to delineate their spatial extent. the three parameters are also calculated using TP, FP, FN for true positive, false positive and false negative spatially connected clusters. A predicted cluster is considered to be a true cluster if it spatially overlaps with a cluster in the ground truth mask. For PA segmentation, the three parameters were calculated both before and after applying WM masks to removed FP voxels.

#### Threshold based PA segmentation

For comparison, we also obtained PA masks using a threshold-based method as described in [10]. Briefly, voxels with signal intensity two standard deviations above the smoothed neighboring intensity were identified in the phase images and the magnitude images with flow encoding gradient off. The PA mask was then defined as clusters in the phase image that overlapped with a cluster in the magnitude image within WM. Clusters within regions with ghosting artifacts due to large surface vessels were manually excluded.

### 2.4 Data analysis

#### 2.4.1 Image reconstruction

The FatNav images were reconstructed using the standard GRAPPA reconstruction algorithm after inverse Fourier transform (IFT) of the data along the readout direction [21]. A separate kernel with a size of 2×2 was estimated for each position along the readout to calculate the missing k-space data. The missing data due to partial Fourier acquisition was zero filled before IFT along PAR and PE directions. The images from all channels were combined into a single image by root mean square. The TSE images were reconstructed by the vendor provided software on the scanner.

PC MRI images were reconstructed offline to a voxel size of 0.1563× 0.1563 mm^2^ by zero padding in k-space before inverse Fourier transform. Images from repetitions 2 – 10 were averaged. The first repetition was discarded because there was inherent slice position inconsistency between k-space data received before and after receiving the first feedback during the first repetition. The inconsistency was caused by subtle differences between the reference FatNav images acquired during the TSE and PC MRI sequences, which caused systematic errors (nonzero biases) in estimated motion parameters. Then, coil-combined phase and magnitude images were obtained as described in [10].

To study whether replacing data affected by motion can further improve image quality, we further replaced data block *D*_i_ in the initial acquisition by a reacquired block *D*_j_ at the same k-space locations if the sum of the maximums of *T*_i-1_, *T*_i_, and *T*_i+1_ (Tmax_i_) and of *R*_i-1_, *R*_i_, and *R*_i+1_ (Rmax_i_) was greater than Tmax_j_+Rmax_j_, before the above image reconstruction.

#### 2.4.2 PA segmentation

Out of the ten trained models for each network, the one with the largest DSC (DSC of WM in the case of U-NET) was selected to segment the PC MRI and TSE images in Exp 2. To generate the WM masks, we first resampled the WM masks generated by U-NET at the pixel positions of the

PC MRI images. Then, the resampled WM masks were manually adjusted to account for possible head movement between the T2w and PC MRI scans (Exp 1) or for above-mentioned systematic error in FatNav image registration (Exp 2). In Exp 1, the WM masks were only available for 50 of the 61 PC MRI images. The number of PAs (N_PA_) is defined as the number of spatially connected clusters in the final masks.

#### 2.4.3 Motion parameters

To investigate whether the N_PA_ difference between MC and NoMC scans depended on the motion severity, rotational (*M*_R_) and translational (M_T_) motion ranges were calculated for all scans. *M*_R_ (*M*_T_) was defined as the root mean square of the differences of the maximum and minimum values of the three rigid-body rotational (translational) parameters of all FatNav images acquired during repetitions 2 – 10.

#### 2.4.4. Statistical Analysis

Wilcoxon’s signed rank tests were performed to compare the PPV, SEN, and DSC values of the various PA segmentation methods.

To evaluate whether MC improves PA visualization, the MC and NoMC image pairs were formed by pairing the NoMC and each of the MC scans from the same subject. The image pairs were visually inspected and those showing clear spatial mismatch (likely due to motion during or before the NoMC scans) between the two images were excluded. As a result, images from 5 out of the 14 subjects were excluded, resulting in 26 pairs for comparison. The pairs were divided into three groups based on their difference in motion parameters. A motion score was defined as: score = *M*_T_ + 57.3 mm x *M*_R_, where *M*_T_ and *M*_R_ are in units of mm and radians, respectively. The motion score is similar to that defined in [22] and corresponds to adding to *M*_T_ a shift caused by rotation *M*_R_ of a point on the surface of a sphere with a radius of 57.3 mm. For simplicity, the radius of 57.3 mm is chosen instead of the 64 mm in [22] such that the score is simply equal to the sum of *M*_T_ and *M*_R_ when they are expressed in mm and degrees, respectively. The pairs with motion score difference between MC and NoMC in the ranges of (-∞, *θ*_M_), [-*θ*_M_, *θ*_M_], and (*θ*_M_, ∞) are defined as having less, similar, and more motion during MC scan, respectively. Similarly, the pairs with V_PA_ difference in the ranges of (-∞, −*θ*_N_), [-*θ*_N_ *θ*_N_], and (*θ*_N_, ∞) are defining as having decreased, similar, and increased *N*_PA_ during MC. In this study, the motion score threshold *Θ*_M_ was set to 0.4 mm which is roughly equal to twice the sum of root mean squares of residual standard deviations of the rotational and translational parameters as defined in our previous study [17]. The N_PA_ threshold *θ*_N_ is chosen as 2 which is twice the average number of false positive PA clusters in testing the performance PA segmentation with M2EDN plus WM masks in Exp 1. Binomial tests were performed to evaluate whether the differences between the numbers of pairs with increased and decreased N_PA_ were significant.

To evaluate whether replacement with reacquired data further improved image quality, signed rank tests were performed to compare N_PA_ between images with and without replacement. Spearman’s correlation coefficients were calculated between the fraction of replaced data and the relative N_PA_ changes (ΔN_PA_) between images with and without data replacement. Relative ΔN_PA_ (in units of %) is defined as 200(*N*_PA,w_ – *N*_PA,wo_)/(*N*_PA,w_ + *N*_PA,wo_), where the subscripts *w* and *wo* denote with and without replacement, respectively. For all statistical tests, p values less than 0.05 are considered significant.

### 2.5. Code and data availability

All images and the trained CNN models will be shared upon publication of the manuscript.

## 3. RESULTS

Table 1 consists of SEN, PPV, and DSC values for the 4 ROIs in our U-NET segmentation. SEN values are highest for thalamus (0.82) and lowest for midbrain (0.68). PPV values across all 4 ROIs range between 0.8-0.89. DSC values range from 0.74 to 0.85 across the 4 ROIs, indicating high accuracy of our U-NET model in its segmentation of the 4 ROIs. Figure 2 shows representative T2w image slices that intersect the four brain regions and the overlaying ROIs generated by the trained U-NET model, demonstrating accurate masks for all ROIs.

**Figure 2:**
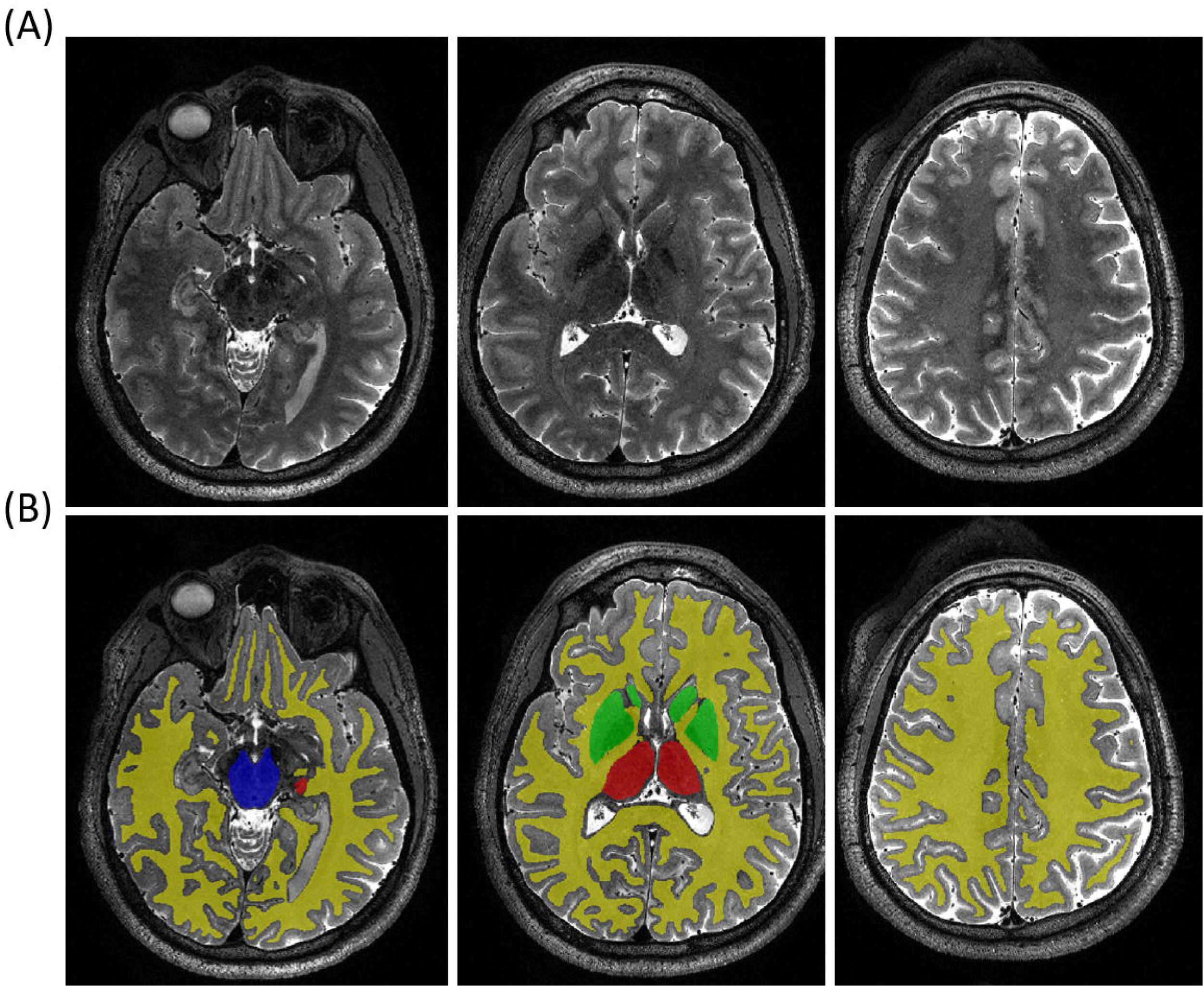
(A) Representative T2w images in a single subject. (B) Overlaid tissue segmentation masks. The yellow, green, red, and blue regions are WM, BG, TH, and MB, respectively.

Figure 3A and 3B show representative PC MRI images and overlaying PA masks, respectively, generated by the M2EDN. Two false PAs were detected in a sulcus of the brain, as denoted by the arrows, which can be removed by applying the WM mask as shown in Fig. 3C. Figure 4 compares various PA segmentation methods side by side. The mask produced by the M2EDN model located four penetrating arteries that the threshold method was unable to identify, as indicated by the 4 green circles. However, the M2EDN model also identifies one false PA as labeled by the blue circle.

**Figure 3:**
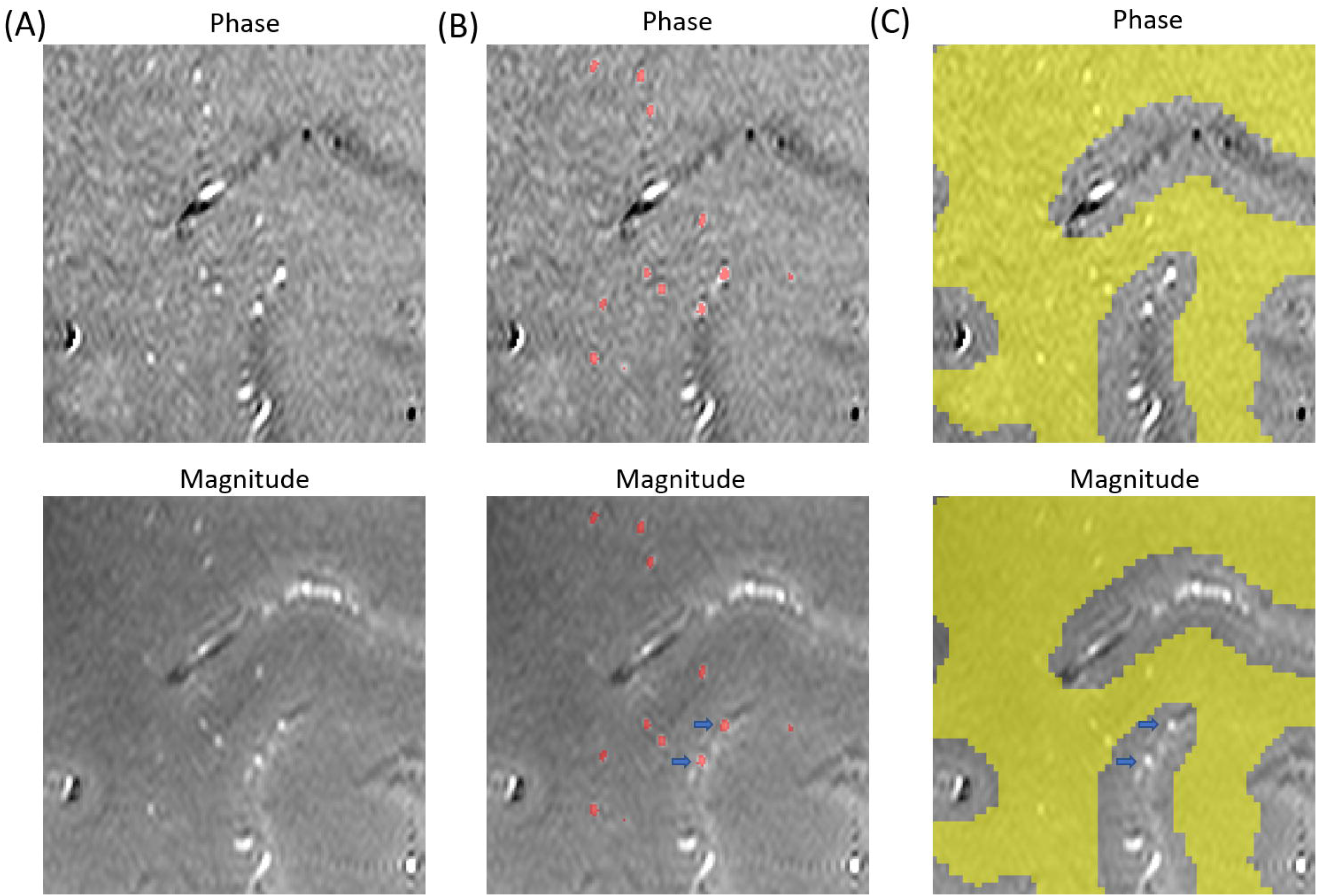
(A) Phase and magnitude images of a representative subject. (B) PA masks generated from the M2EDN network shown as red dots. (C) Resampled WM mask generated from the U-NET shown in yellow. The arrows denote two false positive clusters on the cortical surface.

**Figure 4:**
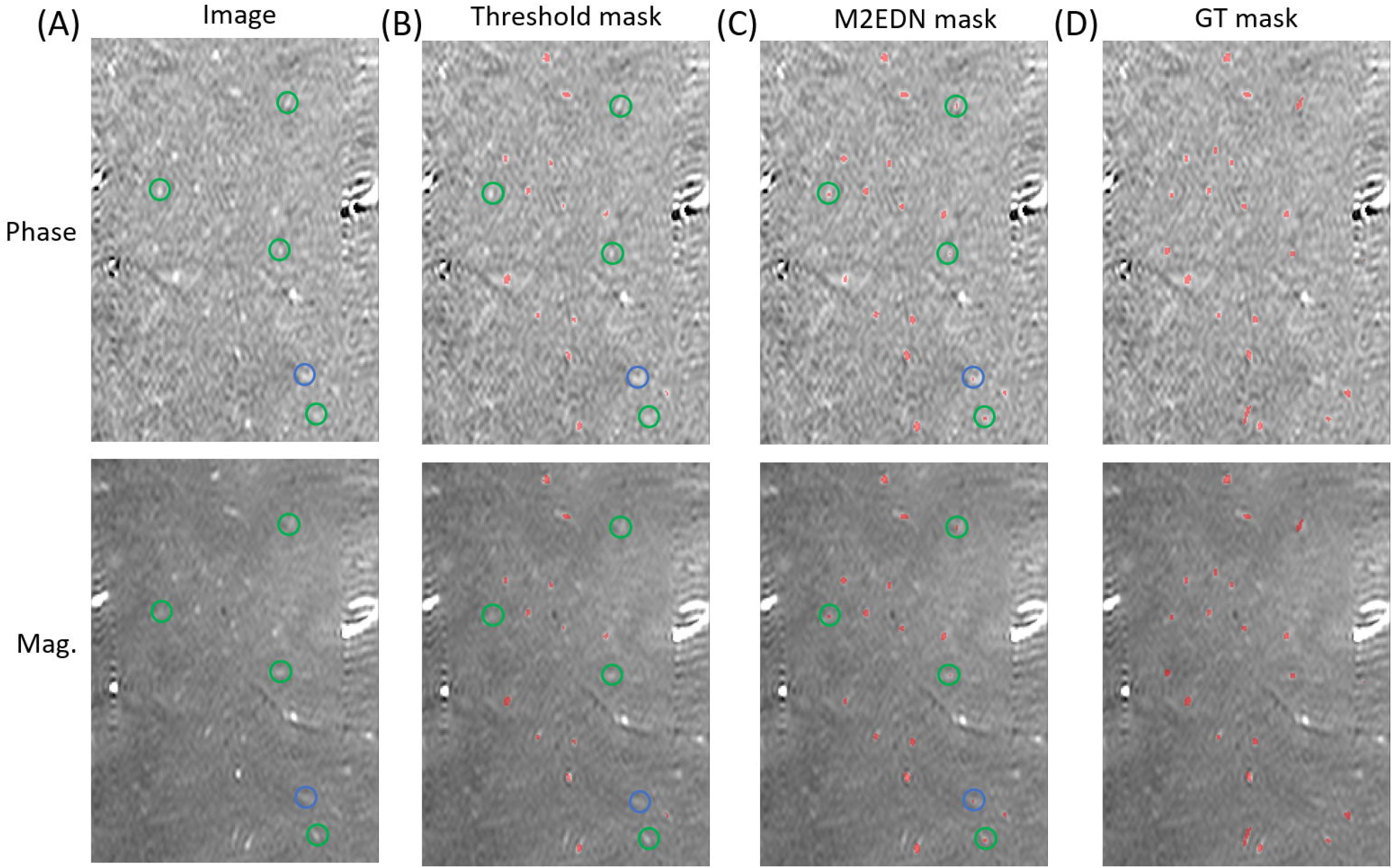
(A) Phase and magnitude images in a representative subject. (B) PA masks generated from the thresholding method, as shown in red. (C) PA masks generated by the M2EDN network as shown in red. (D) the ground truth mask as shown in red. In (A) – (C), the green circles denote true PA clusters that were only detected using M2EDN. The blue circle denotes a false PA cluster in the M2EDN mask that is absent in the threshold mask.

In Table 2, we assess the performance of the M2EDN model vs. threshold method using the SEN, PPV, and DSC parameters. SEN, PPV, and DSC values are in a lower range for voxel level than for cluster level. Applying WM (denoted as M2EDN+WM) had no effect on SEN as expected but significantly increased both PPV (p = 2.4×10^-8^ for both voxel and cluster levels) and DSC (p = 2.4× 10^-8^ for both). When comparing to threshold+WM, M2EDN+WM had higher SEN (p = 4.8×10^-9^ (voxel levels) and 1.2×10^-9^ (cluster level)) and DSC (p = 1.6×10^-8^ and 1.3×10^-9^), but lower PPV (p =6.7×10^-7^ and 5.8×10^-3^). Importantly, DSC values of M2EDN+WM reached a high value of 0.85 versus only 0.71 of threshold+WM at the cluster level.

Figure 5(A) and (B) show phase and magnitude images acquired with prospective MC and with the conventional PC MRI without the FatNav module, respectively, when there was no head motion in both scans. There is no clear difference between the two sets of images, demonstrating that the FatNav module did not negatively affect image quality.

**Figure 5:**
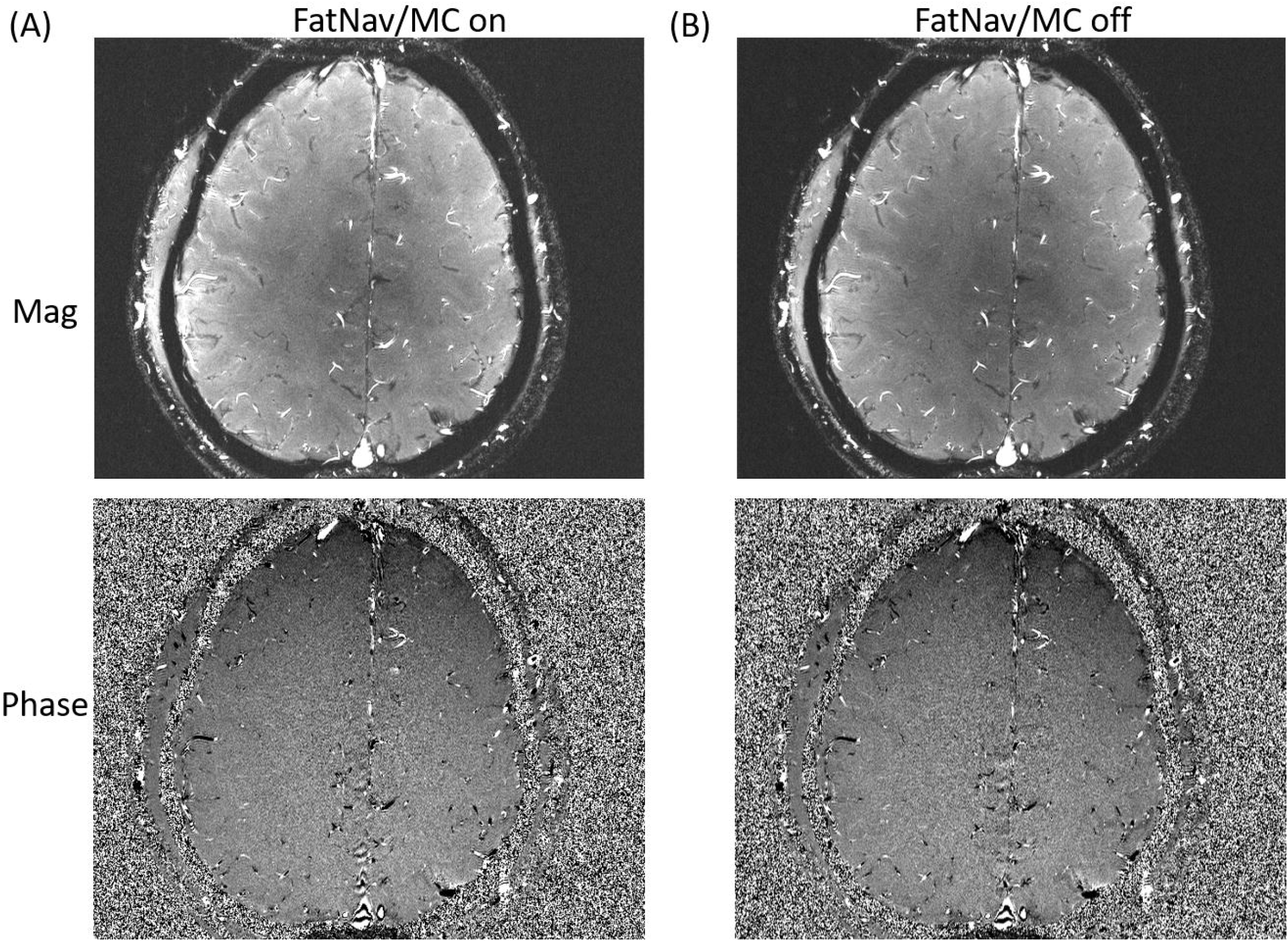
Comparison of PC-MRI images acquired (A) with and (B) without the fat navigator module. The real-time feedback was also turned on in (A).

Table 3 shows a contingency table of cases with various ranges of motion score and NPA differences between MC and NoMC scans. Among the 15 cases where MC and NoMC scans had similar motion scores (second column), MC images showed more NPA in 9 cases (60%) and less N_PA_ in 1 (7%) case (p = 0.011; binomial test). Images for one of the former 9 cases are shown in Figure 6, where PAs can be more clearly visualized on the images with MC but appeared blurry on images with NoMC. As a result, the PAs enclosed by the green circles were only segmented by M2EDN in the MC images. The head motion parameters were *M*_T_ = 0.79 mm (MC) vs 0.59 mm (NoMC) and *M*_R_ = 0.53° (MC) vs 0.54° (NoMC). Motion traces for the two scans can be found in Supporting Information Fig. S2. The motion scores for all the 9 cases are between 0.64 mm and 1.60 mm. In the latter single case where less N_PA_ was obtained with MC, the motion ranges were small (score = 0.56 mm vs 0.64 mm). However, small motion range did not always cause smaller NPA when MC is applied, as the two other MC scans in the same subject also had small motion (score = 0.49 mm and 0.51 mm) but similar N_PA_ as the NoMC scan.

**Figure 6:**
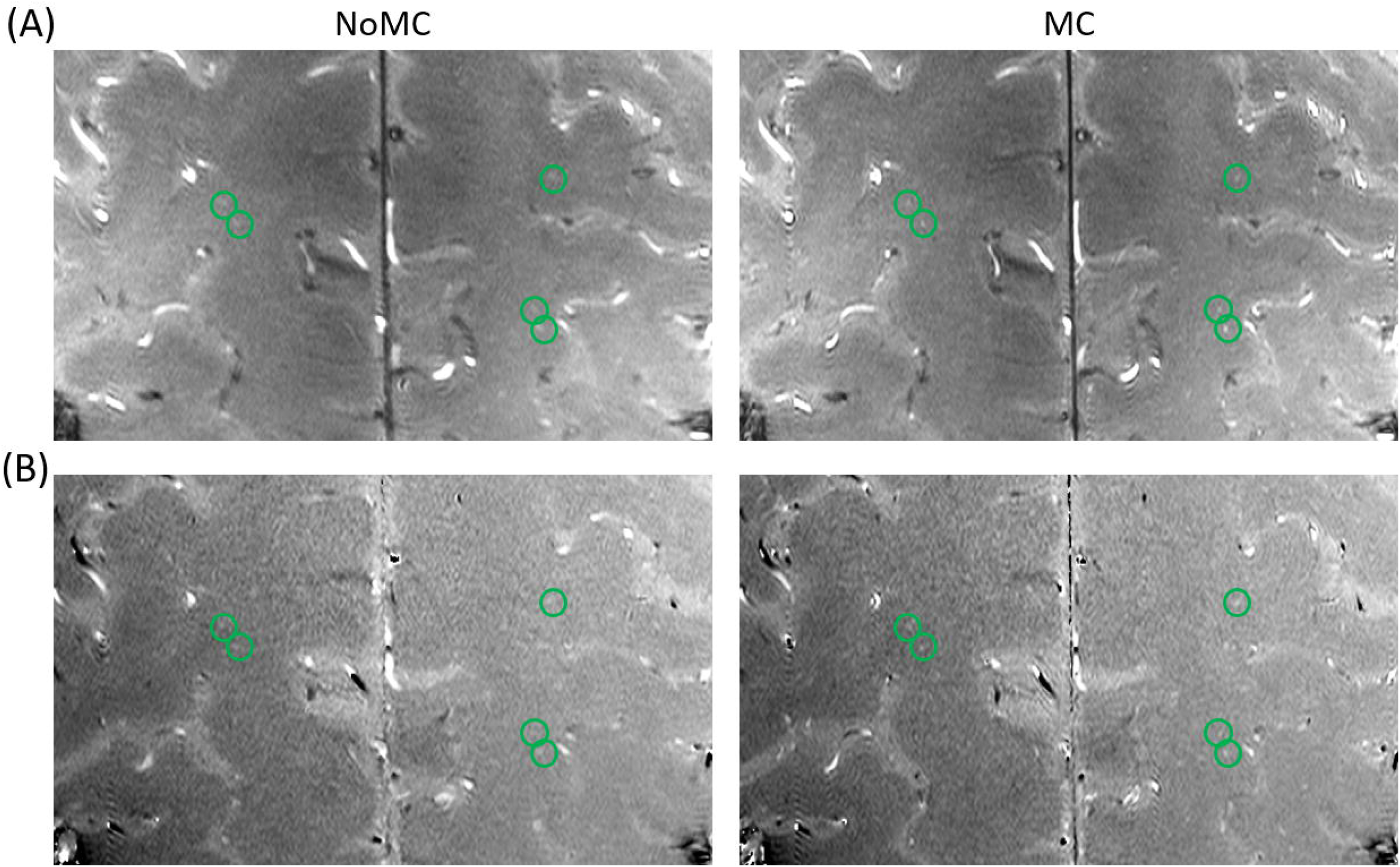
Comparison of (A) magnitude and (B) phase contrast images with MC off (left panel) and on (right panel). The green circles enclose PAs that were only segmented by M2EDN in the MC but not NoMC images.

The third column in Table 3 shows four cases where the MC scans had more motion than the NoMC scans. MC scans had less NPA in three (75%) of the cases, suggesting that MC did not completely eliminate motion artifacts. Those three scans belonged to the same subject, who had the greatest motion (maximum *M*_T_ = 1.2 mm and *M*_R_ = 1.2°) during MC scans and the greatest differences in motion score between the MC and NoMC scans, which equaled to 1.57 mm, 0.83 mm, and 1.3 mm, respectively. In the remaining case, N_PA_ was similar between MC and NoMC scans and the motion score difference was only 0.47 mm. Due to the small number of cases, the difference between cases with less and more N_PA_ (3 vs 0) was not significant (p = 0.125; binomial test). The fourth column shows the distribution of 7 cases where there was less motion during MC than NoMC scans. More and similar N_PA_ were obtained with MC in 6 and 1 cases, respectively. The difference between cases with more and less N_PA_ (6 vs 0) was significant (p = 0.016; binomial test). The list of all *M_T_, M_R_,* N_PA_ values for the MC and NoMC scans can be found on the Supporting Information Table S2.

Figures 7 shows relative ΔN_PA_ versus the fraction of data replaced by reacquired data. No significant correlation (ρ = 0.06; p = 0.64) was observed and signed rank test between N_PA_ values with and without replacement gave a p value of 0.11, suggesting no significant improvement in PA visualization with data reacquisition.

**Figure 7:**
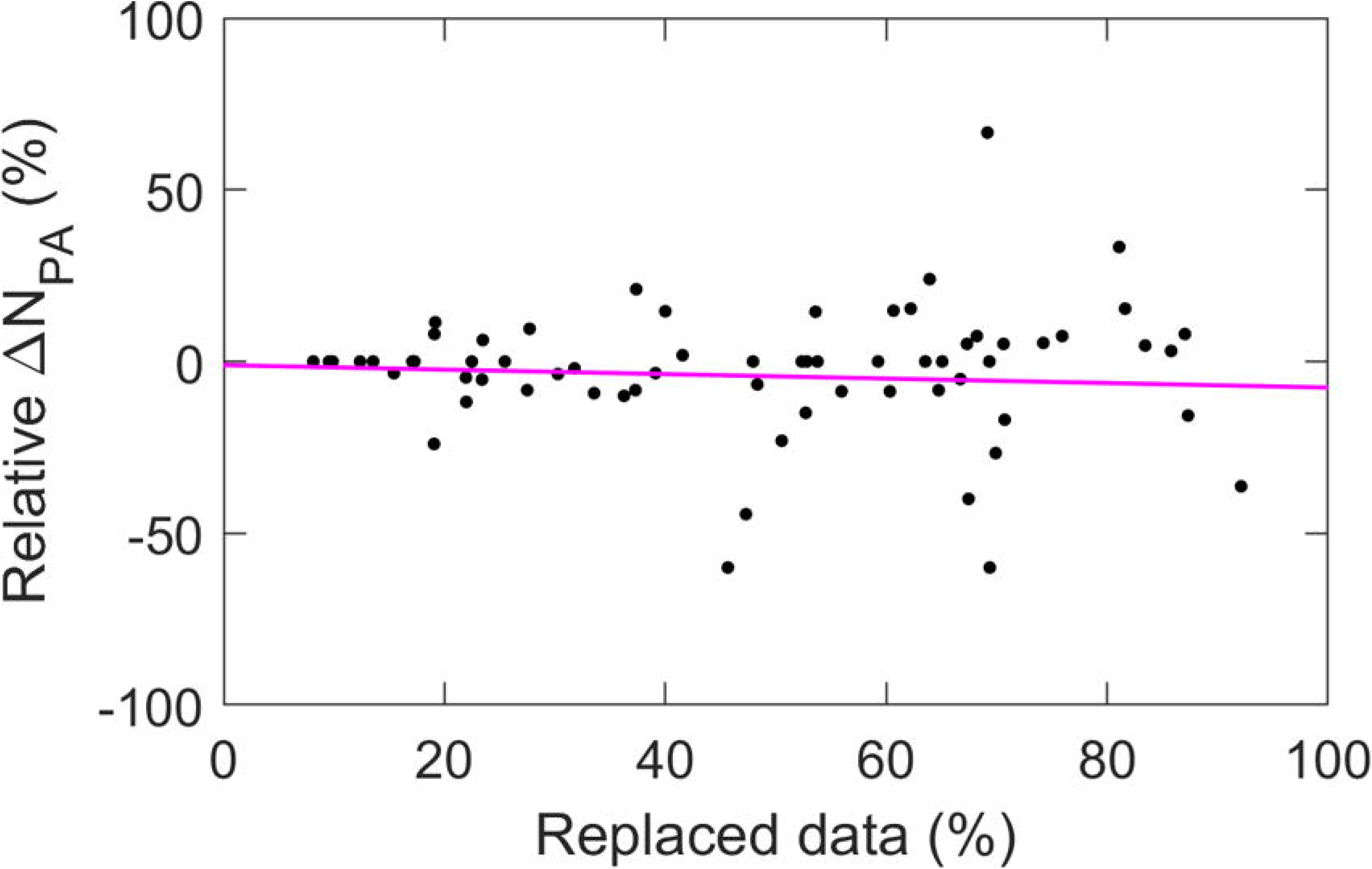
Scatter plots of relative ΔN_PA_ versus the fraction of data replaced by those with smaller motion scores. The red line is a least square fit.

## 4. DISCUSSION

In this study, we developed FatNav based prospective MC method for improving PA visualization in 2D PC MRI sequence and evaluated its performance using automatic PA segmentation methods based on M2EDN and 3D U-NET. Compared to the NoMC scans with similar motion scores, MC significantly improved the visualization of PA and increased the number of segmented PAs. However, we found no further improvement by reacquiring data affected by head motion. The convolutional neural networks increased the SEN and DSC of PA delineation compared to conventional threshold-based method.

Although navigator based prospective MC has been incorporated into various MRI sequences [22–29], it has never been applied to PC MRI. In previous applications, a complete dataset of navigator echo or image were acquired in a single time block. This is impractical for PC MRI due to its short repetition time. Therefore, the k-space data for navigator and main images were acquired in an interleaved fashion, which increased the possibility of artifacts due to the introduction of additional gradient pulses between RF excitations for water signal. However, no visible artifacts were introduced, as illustrated in Fig. 2, which might be explained by the nulling of the zeroth order gradient moment of the FatNav module. A second distinction from previous studies is that the fat signal was used for navigators which avoids reducing the water signal and thus image intensity of the main image.

A shortcoming of the interleaved acquisition is that it decreased the frequency for motion parameter updates, making it challenging to measure and correct sudden motions. We tried to ameliorate this issue by reacquiring data blocks during which motion had occurred. More strict criteria were applied to detect motion than in previous studies by requiring no relative motion between three consecutive FatNav images. The rationale behind the strict criteria is that sudden motion can lead to artifacts in FatNav images which might manifest as changes in motion parameters during image registration. The criteria should be sufficient to detect stepwise motion but its sensitivity for detecting spike-like motion remains unclear.

While improved PA visualization and N_PA_ were observed in scans with MC compared to NoMC, we did not find further improvement with data reacquisition, in apparent contradiction to previous studies where data reacquisition was found to be important for complete correction of image artifacts [22, 25]. There are several possible reasons for the lack of improvement in our study. First, in contrast to the correction of deliberate large motions in the previous studies [22, 25], our study focused on real-life scanning conditions in which subjects were not asked to make deliberate large head movements, but to keep as still as possible. As a result, decent image quality was obtained in most cases even without data reacquisition, leaving less room for further improvement with data reacquisition. Second, as discussed above, sudden spike-like motions may not be detected by FatNav image registration. Third, although ~10 – 90% data blocks were replaced by reacquired data using our criteria, some of them may not truly represent data with more motion due to the intrinsic measurement errors in M_T_ and M_R_ and the small scores of the true motion. A related issue is the slight dissimilarity between the FatNav images and the reference FatNav image acquired during the TSE scans which not only caused systematic bias but might also increase random errors in the estimated motion parameters.

We conducted CNN-based PA and WM segmentation to determine whether the combination of these two masks can provide more accurate segmentation results than the existing threshold based method. According to the data we gathered, the new method identified a greater number of PAs than the previous threshold method. In Table 1, it is evident that the CNN-based method improved overall accuracy of PA identification. PPV values were lower for both M2EDN and M2EDN+WM and SEN values were higher compared to those of threshold+WM. A lower PPV value indicates a higher false positive rate and this is exhibited in Figure 3C as the M2EDN mask identifies one false artery that the threshold method did not identify. SEN values for the M2EDN masks are much higher than the threshold method indicating fewer false negatives. Adding the WM mask on top of the PA segmentation obtained slightly greater DSC values as opposed to M2EDN alone, suggesting that the WM mask was effective in excluding false PAs located on the cortical surface.

Although CNN-based tissue segmentation of 7 T healthy brain MRI images has been reported before [30, 31], our study is the first to evaluate the performance of such an approach for WM segmentation based on T2w images. CEREBRUM-7T employed 3 layers of 3D convolutional blocks and achieved DSC of 0.9 and 0.86 in WM and BG [30], respectively, which are superior to the DSC results in Table 1. However, T_1_ weighted images which have higher tissue contrast were employed. Khandelwal et. al. segmented gray matter in 7 T ex vivo T2w images with 0.3×0.3×0.3 mm3 resolution and achieved DSC of 78.5% to 98.5%. However, the WM was not segmented [31].

There are several proposed causes of SVD. One in particular is the occlusion of PAs which can result in a lacunar infarct downstream of the occlusion. On PC MRI images, occluded arteries without blood flow will not appear on the scan. Fewer identified arteries in our segmentations could be an indicator of potential diseased PAs. Longitudinal studies could assess how other factors such as cholesterol level, blood pressure, or blood glucose levels affect the blood flow through PAs. Observing whether blood flow improves or worsens with therapeutic interventions could be useful for developing effective prevention and treatment strategies for SVD.

Our study has several limitations. First, due to different sequence parameters for FatNav between T2w and PC MRI scans, there existed a systematic error in the motion parameters and as a result the first repetition of PC MRI had to been discarded. In the future, more accurate image registration algorithm should be developed and incorporated into the sequence. Second, in evaluating the performance of MC, controlled deliberate large motion was typically performed. However, we did not perform such comparison, but instead based our evaluation on real-life data, in which the motion is only ≤ 1.2 mm and 1.2 deg. In the subject with the largest motion (Subject 2 in Table S2 and column 3 row 2 in Table 3), MC did not completely overcome the detrimental effects of motion and had less PAs than the NoMC images which had much less motion. However, more cases are needed to further test whether MC can still be effective in cases with large motion (motion score ~ or > 1 mm). Third, due to the relative long TR to acquire the complete k-space data for each FatNav image, spike like motion may not be detected. Other motion detection approaches such as that based on free induction decay signal should be explored in future development [31], which may enable further improvement in image quality by data reacquisition.

## 5. CONCLUSIONS

We have developed a prospective motion correction method for PC MRI which improved the visualization of PA and segmented PA counts in centrum semiovale at 7 T. Furthermore, we have improved the overall accuracy (DSC) and sensitivity of PA segmentation using CNN-base approaches. The developed PA imaging and segmentation methods may help illuminate the mechanisms of pathophysiological changes of PAs in SVD.

## ACKNOWLEDGEMENTS

The project described was partly supported by United States National Institutes of Health through grant 5R21NS095027-02 and the National Center for Advancing Translational Sciences (NCATS), National Institutes of Health, through Grant Award Number UL1TR002489.

Supporting Information Figure S1: The architecture of the 3D U-NET for brain tissue segmentation. The numbers above the rectangles specify the input/output channels of different layers, while the numbers above the pink arrows specify the dropout rate of the Dropout layer.

Supporting Information Figure S2: Motion parameter traces during the (A) NoMC and (B) MC scans in Fig. 6, showing similar degrees of motion during the MC on and off scans. Different colors represent different motion directions or rotation axes.

## TABLE CAPTIONS

Table 1: SEN, PPV, and DSC of TH, BG, MB, and WM masks obtained using UNET segmentation of T2w images.

Table 2: SEN, PPV, and DSC of penetrating artery masks obtained using the M2EDN, M2EDN+WM, and threshold+WM methods. The numbers 61 and 50 inside the parentheses of the second column correspond to all images and those with WM masks, respectively, in Exp. 1. The # and + signs denote significant difference from the corresponding M2EDN and threshold+WM results, respectively.

Table 3: The contingency table of cases with different motion score and N_PA_ relationships between a MC scan and the NoMC scan in the same subject. The labels refer to MC relative to NoMC. The – and + signs in the parentheses denote decline and improvement, respectively, in PA visibility due to MC, as compared to NoMC.

Supporting Information Table S1: MRI parameters for the MPRAGE, VFA-TSE, and MP2RAGE sequences. In the PC sequence, VENC = 4 cm/s and a one-sided flow encoding were employed, thereby the flow encoding gradient was turned on and off alternatively in different TRs.

Supporting Information Table S2: list of the subject ID, NPA, MT, MR, and MT+MR difference for all 26 image pairs grouped according to the NPA and motion range differences between MC and NoMC scans.

## Notes

### Competing Interest Statement

The authors have declared no competing interest.

