## Supplementary figures and images for "Prospective Motion Correction and Automatic Segmentation of Penetrating Arteries in Phase Contrast MRI at 7 T"

### Figure S1

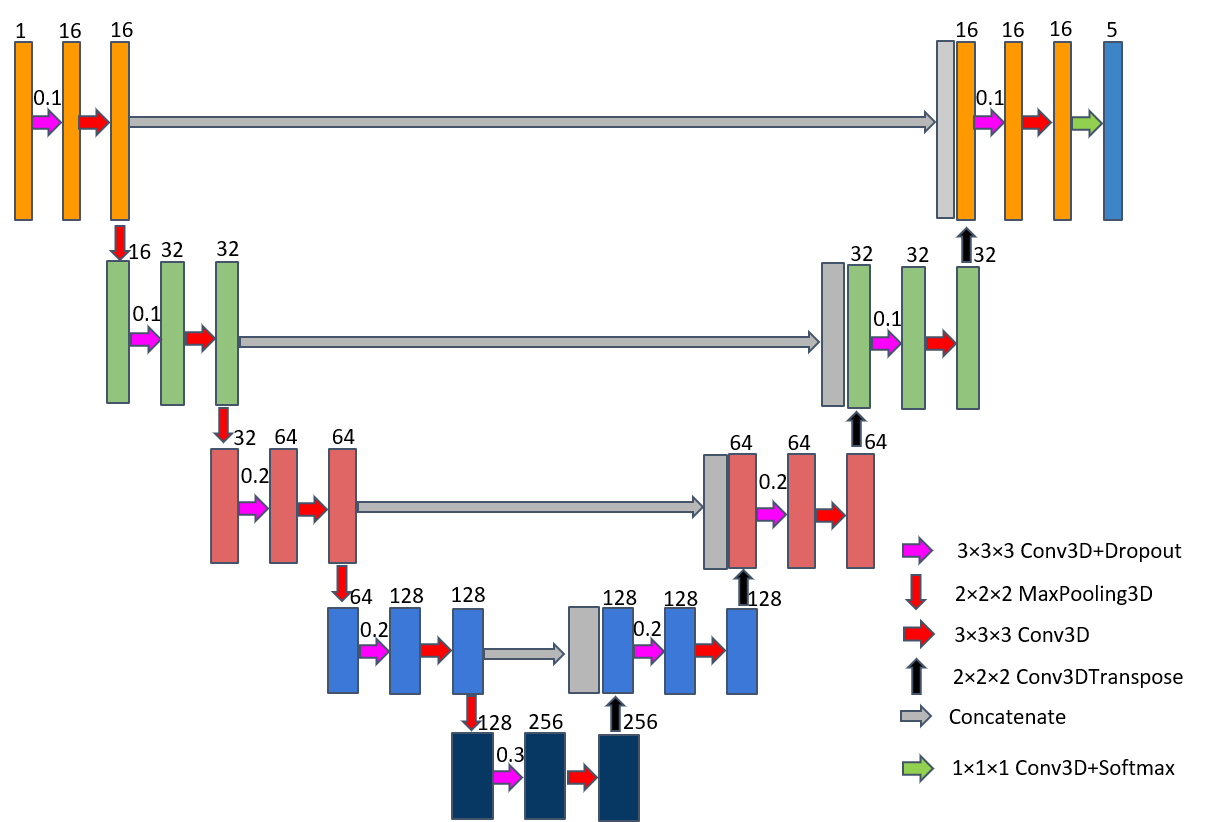

### Figure S2

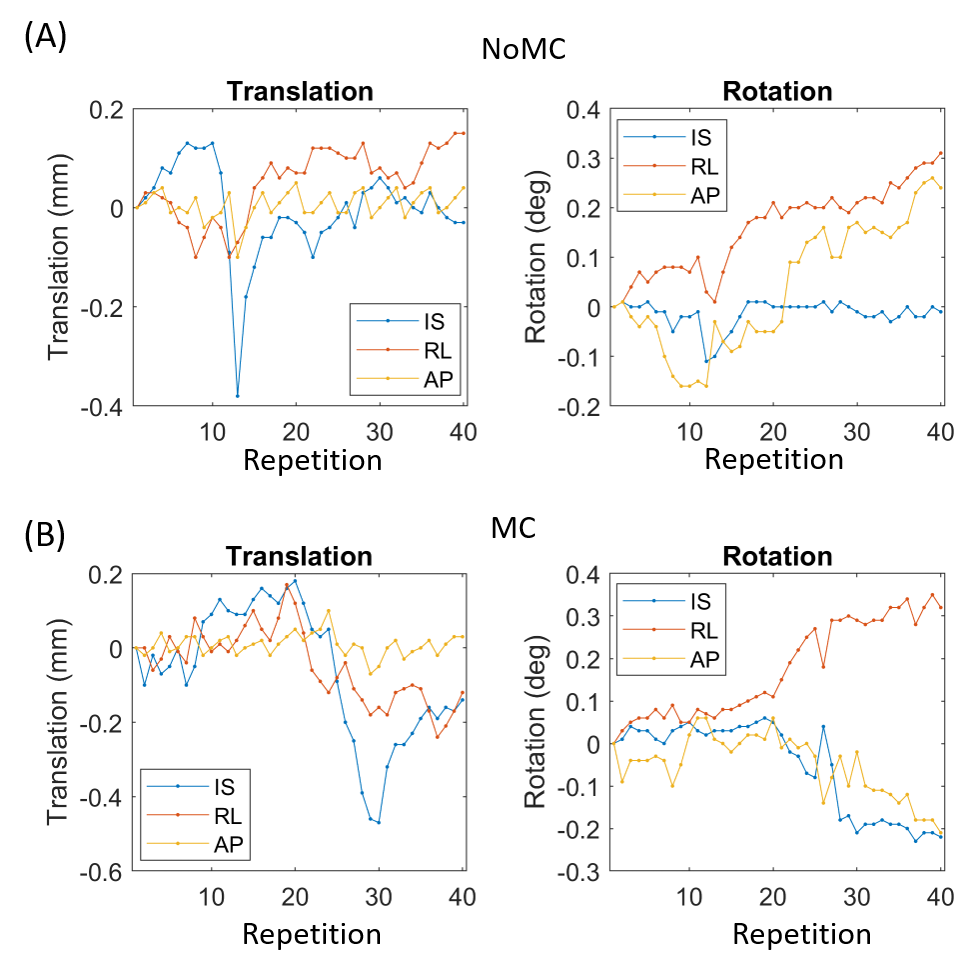
