## Supplementary material for "Prospective Motion Correction and Automatic Segmentation of Penetrating Arteries in Phase Contrast MRI at 7 T": Online supplementary material

|  | TSE | PC | FatNav (TSE in EXP 1) | FatNav (Exp 2) |
| --- | --- | --- | --- | --- |
| TE (ms) | 326 | 15.7 | 1.5 | 1.31 |
| TR (ms) | 3000 (Exp 1)  3300 (Exp 2) | 30 | 3.1 | 30 (PC)  3 (TSE) |
| FOV (mm^3^) | 210×210×99.2 | 200×162.5×2 | 220×220×180 | 222×198×210 |
| Acquired voxel size (mm^3^) | 0.41×0.4×0.4 | 0.31×0.52×2 | 2.2×2.2×2.2 | 3×3×3 |
| Recon voxel Size (mm^3^) | 0.41×0.41×0.4 | 0.156×0.156×2 | 2.2×2.2×2.2 | 3×3×3 |
| Recon matrix Size | 512×512×248 | 1280×768×1 | 100×100×82 | 74×66×70 |
| FA (degree) | Variable | 45 | 7 | 14 (PC)  7 (TSE) |
| Slice Orientation | Axial | Oblique | Axial | Axial |
| Partial Fourier Factor (PE×PAR) | 0.79×0.625 | 1×1 | 0.75×0.75 | 0.75×0.75 |
| GRAPPA factor (PE×PAR) | 3×1 | 1×1 | 4×4 | 4×4 |
| GRAPPA ACS lines | 24 | NA | 32×20 | 21×23 |
| Bandwidth (Hz/Pixel) | 349 |  |  |  |
| TA | 8:03 min | 3:13 min (10 repetitions) | 0.89 s (Exp 1) | 4.68 s (PC)  0.47 s (TSE) |

Supporting Information Table S1: MRI parameters for the MPRAGE, VFA-TSE, and MP2RAGE sequences. In the PC sequence, VENC = 4 cm/s and a one-sided flow encoding were employed, thereby the flow encoding gradient was turned on and off alternatively in different TRs.

|  | Sub.  ID | N_PA_ | | M_T_ | | M_R_ | | M_T_+M_R_ difference |
| --- | --- | --- | --- | --- | --- | --- | --- | --- |
|  |  | MC | NoMC | MC | NoMC | MC | NoMC |  |
| Less N_PA_  Similar motion | 9 | 28 | 33 | 0.229 | 0.299 | 0.332 | 0.340 | -0.078 |
| Less N_PA_  More motion | 2 | 20 | 49 | 1.239 | 0.302 | 1.047 | 0.416 | 1.567 |
|  | 2 | 20 | 49 | 0.686 | 0.302 | 0.865 | 0.416 | 0.832 |
|  | 2 | 25 | 49 | 0.848 | 0.302 | 1.153 | 0.416 | 1.282 |
| Similar N_PA_  Less motion | 3 | 32 | 31 | 0.488 | 0.649 | 0.287 | 0.536 | -0.410 |
| Similar N_PA_  Similar motion | 6 | 22 | 21 | 0.492 | 0.273 | 0.330 | 0.283 | 0.266 |
|  | 9 | 32 | 33 | 0.215 | 0.299 | 0.273 | 0.340 | -0.151 |
|  | 3 | 29 | 31 | 0.498 | 0.649 | 0.341 | 0.536 | -0.345 |
|  | 6 | 21 | 21 | 0.380 | 0.273 | 0.540 | 0.283 | 0.364 |
|  | 9 | 34 | 33 | 0.263 | 0.299 | 0.245 | 0.340 | -0.131 |
| Similar N_PA_  More motion | 4 | 13 | 14 | 0.670 | 0.315 | 0.507 | 0.396 | 0.466 |
| More N_PA_  Less motion | 3 | 41 | 31 | 0.375 | 0.649 | 0.271 | 0.536 | -0.539 |
|  | 8 | 41 | 37 | 0.241 | 0.536 | 0.520 | 1.070 | -0.846 |
|  | 5 | 24 | 9 | 0.254 | 0.380 | 0.481 | 0.975 | -0.621 |
|  | 8 | 44 | 37 | 0.234 | 0.536 | 0.604 | 1.070 | -0.768 |
|  | 5 | 19 | 9 | 0.292 | 0.380 | 0.582 | 0.975 | -0.482 |
|  | 8 | 50 | 37 | 0.313 | 0.536 | 0.467 | 1.070 | -0.826 |
| More N_PA_  Similar motion | 1 | 13 | 0 | 0.601 | 0.587 | 0.388 | 0.536 | -0.134 |
|  | 5 | 13 | 9 | 0.461 | 0.380 | 0.535 | 0.975 | -0.359 |
|  | 7 | 28 | 23 | 0.819 | 0.928 | 0.441 | 0.649 | -0.317 |
|  | 1 | 10 | 0 | 0.787 | 0.587 | 0.529 | 0.536 | 0.193 |
|  | 4 | 19 | 14 | 0.370 | 0.315 | 0.357 | 0.396 | 0.016 |
|  | 7 | 29 | 23 | 0.983 | 0.928 | 0.618 | 0.649 | 0.024 |
|  | 1 | 11 | 0 | 0.477 | 0.587 | 0.419 | 0.536 | -0.227 |
|  | 4 | 19 | 14 | 0.314 | 0.315 | 0.326 | 0.396 | -0.071 |
|  | 7 | 29 | 23 | 0.774 | 0.928 | 0.424 | 0.649 | -0.379 |

Supporting Information Table S2: list of the subject ID, NPA, MT, MR, and MT+MR difference for all 26 image pairs grouped according to the NPA and motion range differences between MC and NoMC scans.
