## Supplementary material for "Prospective Motion Correction and Automatic Segmentation of Penetrating Arteries in Phase Contrast MRI at 7 T": Tables

|  | SEN | PPV | DSC |
| --- | --- | --- | --- |
| TH | 0.82 (0.14) | 0.89 (0.059) | 0.85 (0.09) |
| BG | 0.8 (0.091) | 0.8 (0.15) | 0.79 (0.074) |
| MB | 0.68 (0.19) | 0.86 (0.051) | 0.74 (0.15) |
| WM | 0.77 (0.095) | 0.86 (0.081) | 0.8 (0.043) |

|  |  | SEN | PPV | DSC |
| --- | --- | --- | --- | --- |
| Voxel level | M2EDN (n=61) | 0.57 (0.13) | 0.79(0.07) | 0.65(0.09) |
|  | M2EDN (n=50) | 0.56 (0.12) | 0.80 (0.07) | 0.64(0.08) |
|  | M2EDN+WM (n=50) | 0.56 (0.12)^+^ | 0.84 (0.06) ^#,+^ | 0.66 (0.09)^#,+^ |
|  | threshold + WM (n=50) | 0.38 (0.13) | 0.92 (0.10) | 0.52 (0.14) |
| Cluster level | M2EDN (n=61) | 0.84(0.10) | 0.82(0.10) | 0.82(0.06) |
|  | M2EDN (n=50) | 0.83 (0.10) | 0.82 (0.09) | 0.82 (0.06) |
|  | M2EDN+WM (n=50) | 0.83 (0.10)^+^ | 0.88 (0.08)^#,+^ | 0.85 (0.06)^#,+^ |
|  | threshold + WM (n=50) | 0.59 (0.13) | 0.92 (0.08) | 0.71 (0.10) |

Table 1: SEN, PPV, and DSC of TH, BG, MB, and WM masks obtained using UNET segmentation of T2w images.

Table 2: SEN, PPV, and DSC of penetrating artery masks obtained using the M2EDN, M2EDN+WM, and threshold+WM methods. The numbers 61 and 50 inside the parentheses of the second column correspond to all images and those with WM masks, respectively, in Exp. 1. The # and + signs denote significant difference from the corresponding M2EDN and threshold+WM results, respectively.

|  | Similar motion | More motion | Less motion |
| --- | --- | --- | --- |
| Less N_PA_ | 1 | 3 | 0 |
| Similar N_PA_ | 5 | 1 | 1 |
| More N_PA_ | 9 | 0 | 6 |

Table 3: The contingency table of cases with different motion score and N_PA_ relationships between a MC scan and the NoMC scan in the same subject. The labels refer to MC relative to NoMC. The – and + signs in the parentheses denote decline and improvement, respectively, in PA visibility due to MC, as compared to NoMC.
